# Circular RNAs regulate cancer stem cells by FMRP against CCAR1 complex in hepatocellular carcinoma

**DOI:** 10.1101/396069

**Authors:** Yan-Jing Zhu, Bo Zheng, Xu-Kai Ma, Xin-Yuan Lu, Xi-Meng Lin, Gui-Juan Luo, Shuai Yang, Qing Zhao, Xin Chen, Ying-Cheng Yang, Xiao-Long Liu, Rui Wu, Jing-Feng Liu, Yang Ge, Li Yang, Hong-Yang Wang, Lei Chen

**Author notes:** **Corresponding Authors: Lei Chen**, Ph.D. International Co-operation Laboratory on Signal Transduction, Eastern Hepatobiliary Surgery Institute, Second Military Medical University, 225 Changhai Road, 200438 Shanghai, China. **Hong-Yang Wang**, M.D., Ph.D. International Co-operation Laboratory on Signal Transduction, Eastern Hepatobiliary Surgery Institute, Second Military Medical University, 225 Changhai Road, 200438 Shanghai, China. These authors contributed equally to this work (Yan-Jing Zhu, Bo Zheng, Xu-Kai Ma, Xin-Yuan Lu).

## Abstract

Circular RNA (circRNA) possesses great pre-clinical diagnostic and therapeutic potentials in multiple cancers. However, the underlying correlation between circRNAs and cancer stem cells (CSCs) has not been reported. The absence of circZKSCAN1 endowed several malignant properties including cancer stemness and tightly correlated with worse overall and recurrence-free survival rate in HCC cells in vitro and in vivo. Bioinformatics analysis and RNA immunoprecipitation-sequencing (RIP-seq) results revealed that circZKSCAN1 exerted its inhibitive role by competitively binding FMRP, therefore, block the binding between FMRP and β-catenin-binding protein-cell cycle and apoptosis regulator 1 (CCAR1) mRNA, and subsequently restraining the transcriptional activity of Wnt signaling. In addition, RNA-splicing protein Quaking 5 was found downregulated in HCC tissues and responsible for the reduction of circZKSCAN1. Collectively, this study revealed the mechanisms underlying the regulatory role of circZKSCAN1 in HCC CSCs and identified the newly discovered Qki5– circZKSCAN1–FMRP–CCAR1–Wnt signaling axis as a potentially important therapeutic target for HCC treatment.

**Statement of significance:**

1. CircZKSCAN1, a novel negative regulator for cancer stem cells, was firstly identified with reverse correlation with HCC prognosis.
2. CircZKSCAN1 directly targets FMRP, and competitive binding with β-catenin-binding protein cell cycle and apoptosis regulator 1 (CCAR1) for its activity.
3. The decreased expression of Quaking 5 is responsible for the absence of circZKSCAN1 in HCC.

## Introduction

Hepatocellular carcinoma (HCC) is one of the most lethal malignancies worldwide, accounting for approximately 700,000 deaths per year (1). Liver cancer stem cells (CSC) are defined by stemness-related markers including CD133 and epithelial cell adhesion molecule (EpCAM) and are considered to be responsible for the initiation of tumor progression, drugresistance, metastasis, and recurrence (2). Numerous long noncoding RNAs (lncRNAs) and microRNAs (miRNAs) were found to be dysregulated in HCC, with pro-or anti-tumor properties. LncTLSNC8 and miRNA-708 acted as tumor suppressors in HCC (3,4), while lncBRM and miRNA-429 showed tumor-promoting properties (5,6). However, few studies have examined the involvement of non-coding RNAs, especially circular RNAs (circRNAs) in the regulation of liver CSCs.

CircRNAs are considered to be rare, compared with linear mRNAs with 5′ and 3′ terminal structures reflecting the start and stop signals for transcription of their corresponding DNA templates. The biogenesis of circRNAs is mostly through the back-splicing mechanism. Transcriptome data for humans, mice, and nematodes suggests the existence of large numbers of circRNAs with unidentified functions (7). Furthermore, circRNAs are stable, conserved, and resistant to exonucleolytic RNA decay compared with their source mRNA. CircRNAs feature cell-type-specific and developmental-stage-specific expression, indicating their broad participation in various physiological and pathophysiological processes (7,8).

Generally, circRNAs often possess multiple miRNA-targeting sites and act as endogenous competing RNAs or miRNA sponges to regulate transcriptional activity (9). FOXO3-derived circRNAs prevent tumor growth and metastasis in breast cancer, via performing a ‘sponge’ function involving eight different miRNAs (10). A recent study revealed that circMTO1 acted as a tumor suppressor by binding miRNA-9, which upregulated the expression of p21 in HCC (11).

Till now, it is still unclear whether circRNAs could exert their potential activities during tumor progression beyond miRNA or mRNA sponge. RNA-binding proteins (RBPs), play a crucial role in post-transcriptional modification and mRNA translation. However, whether circRNAs could modulate the malignant behavior of tumors through targeting RBPs has not been documented previously.

Zinc-finger protein with KRAB and SCAN domains 1 (ZKSCAN1) is a member of the zinc-finger protein family, which are classic regulators of gene transcription and closely related to tumorigenesis, tumor progression, and metastasis. ZKSCAN1 expression was shown to be upregulated in esophageal adenocarcinoma and facilitated the proliferation of tumor cells (12). Transcriptome sequencing showed exon–exon junction reads connecting the 5′ end of exon 2 and 3′ end of exon 3, suggesting the possible existence of circZKSCAN1 (circBase ID: hsa_circ_0001727) (8). RNA-seq revealed 372 different gene expression patterns in the circZKSCAN1-knockdown versus control group, while KEGG enrichment analysis suggested that most differentially expressed genes were located in the migration, adhesion, actin cytoskeleton, cytokine interaction, and phosphatidylinositide 3-kinase pathways (13). However, the mechanisms whereby circZKSCAN1 affects the expression of key genes in specific pathways, and whether its inhibitory role in HCC is related to its regulatory effects on miRNAs and RBPs remains unclear.

In the present study, we demonstrated that circZKSCAN1 suppressed cell stemness in HCC through regulating the function of the RBP fragile X mental retardation protein (FMRP). We also identified the downstream target gene of FMRP as cell cycle and apoptosis regulator 1 (CCAR1), which acts as a co-activator of the Wnt/β-catenin signaling pathway and upregulates cell stemness. The results of this study revealed a novel mechanism whereby circRNAs regulate malignant behavior in cancers, in addition to acting as a miRNA sponge. This information may provide a new angle for further research into the relationship between circRNAs and cancers.

## Materials and Methods

### Patients and tissue specimens

In total, 112 pairs of HCC and paired paracancerous tissues were obtained from surgical resections of patients without preoperative treatment at Eastern Hepatobiliary Surgery Hospital (Shanghai, China) from 2009-2016. The procedure of human specimen collection was approved by the Ethics Committee of Eastern Hepatobiliary Surgery Hospital. 20 fresh HCC specimens were obtained from patients undergoing hepatectomy at Eastern Hepatobiliary Surgery Hospital.

All 20 samples were received in the laboratory within 20 min followed by immediately mechanically disaggregated, digested and isolated for further experiments.

### RIP sequencing and bioinformatics analysis

The RIP procedure was performed with Magna RIP Kit (Catalog No. 17-700, Millipore). HCC cell lines were lysed in RIP lysis buffer and then immunoprecipitated with antibody to RBP of interest with protein A/G magnetic beads. Magnets were used to immobilize magnetic beads bound complexes while washing off the unbound materials. Subsequently, remaining RNA was extracted.

Total FMRP-RIP-RNA from hepatocellular carcinoma cell lines was purified with RNeasy Micro Kit according to manufacturer’s instructions (Qiagen, Hilden, Germany).TruSeq Stranded Total RNA with Ribo-Zero kit (Illumina, San Diego, CA) has been used for library preparation following the manufacturer’s instructions. Both RNA samples and final libraries were quantified by using the Qubit 2.0 Fluorometer(Invitrogen, Carlsbad, CA) and quality tested by Agilent 2100 Bioanalyzer RNA Nano assay (Agilent Technologies, Santa Clara, CA). Libraries were then processed with Illumina cBot for cluster generation on the flowcell, following the manufacturer’s instructions and sequenced on 150 bp pair-ends mode at the on HiSeq2500 (Illumina, San Diego, CA). The library construction and sequencing were performed at Shanghai Biotechnology Corporation. Raw sequence files were subjected to quality control analysis using FastQC (https://www.bioinformatics.babraham.ac.uk/projects/fastqc/). Sequencing raw reads were preprocessed by filtering out rRNA reads, sequencing adapters, short-fragment reads and other low-quality reads. We used Tophat v2.0.9 to map the cleaned reads to the human reference genome ensemble GRCh38 (hg38) with two mismatches. After genome mapping, Cufflinks v2.1.1 was run with a reference annotation to generate FPKM values for known gene models. Differentially expressed genes were identified using Cuffdiff. The p-value significance threshold in multiple tests was set by the false discovery rate (FDR). The fold-changes were also estimated according to the FPKM in each sample. The differentially expressed genes were selected using the following filter criteria: FDR ≤0.05 and fold-change ≥2 in hepatocellular carcinoma cell lines. For further analysis, we selected effective data using FMRP IP is up-regulation with IgG group. Complete RIP-seq datasets can be accessed through GEO via series GSE110048.

### Fluorescence in situ hybridization

Fluorescence in situ hybridization was conducted in HCC cell line with CY3-labelled RNA fluorescence probes (GENESEED) against circZKSCAN1 and miR-CCAR1. SMMC7721 cells were grown to 50–75% confluence at the time of fixation. After prehybridization, the cells were hybridized to CY3-labeled probes specific to circZKSCAN1 or CCAR1 at 37 °C overnight. Signals were detected under a microscope (Olympus Corp.). Primers for FISH analysis were Has circ cZKSCAN1-FISH probe: TCTTACAGTCACGAG/GAATAGTAAAGAAAC, Has-miR-CCAR1 (Gene ID: 55749, NM_018237.3).

### In situ hybridization

In situ RNA hybridization was conducted in formaldehyde-fixed liver tissues with digoxygenin (DIG)-labelled RNA probes (Exiqon) against circZKSCAN1 and sense negative control. Anti-DIGAP (Bosterbio, USA) was used as secondary antibody and staining was developed with NBT/BCIP solution (Bosterbio, USA).

## Results

### 1. CircRNA expression spectrum in HCC

The expression patterns of miRNAs and lncRNAs in HCC have been well studied and their diagnostic and therapeutic potentials have been noted; however, the possible dysregulation of circRNAs in HCC remains unknown. We extracted RNA (excluding rRNA) from ten HCC samples and paired paracancerous tissues and performed second-generation sequencing to detected dysregulated RNAs and circRNA patterns in HCC (Fig. 1a and Supplementary Table 1). Gene-cluster analysis showed that the expression levels of numerous genes were dysregulated, with most genes being upregulated in the context of HCC (Fig. 1b and Supplementary Fig. 1a). GO enrichment analysis revealed that these dysregulated genes participated in a wide range of crucial cellular activities (Supplementary Fig. 1b). CircRNA cluster analysis indicated that dysregulated circRNAs were not preferentially up-or downregulated in HCC. However, after circRNA/gene standardization, most circRNAs were shown to be downregulated in HCC (Fig. 1c and d). Based on data from the normal paracancerous tissues, we identified 3,198 liver-specific circRNAs not expressed in other tissues (Fig. 1e and Supplementary Table 2). Considering the blank left demonstrating the relationship between CSCs and circRNAs, we separated the 10 tumor tissues into EpCAM^high^(n = 6) and EpCAM^low^(n = 4) subgroups based on the RNA-seq results (Supplementary Fig. 1c and Supplementary Table 1) and immunohistochemical staining scores (Fig. 1f). The sequencing data were then reorganized based on the newly defined subgroups (Supplementary Fig. 1d and e). Finally, we identified 18 circRNAs that met the following criteria: 1, target circRNA > 2-fold downregulated in HCC; or 2, target circRNA > 1.5-fold upregulated in EpCAM^low^subgroup (Fig. 1g). Among the 18 identified circRNAs, circZKSCAN1 was previously shown to play a distinctive role in HCC, though its mechanism is unknown (13). We, therefore, chose to investigate circZKSCAN1 as a target circRNA.

**Table 1:**
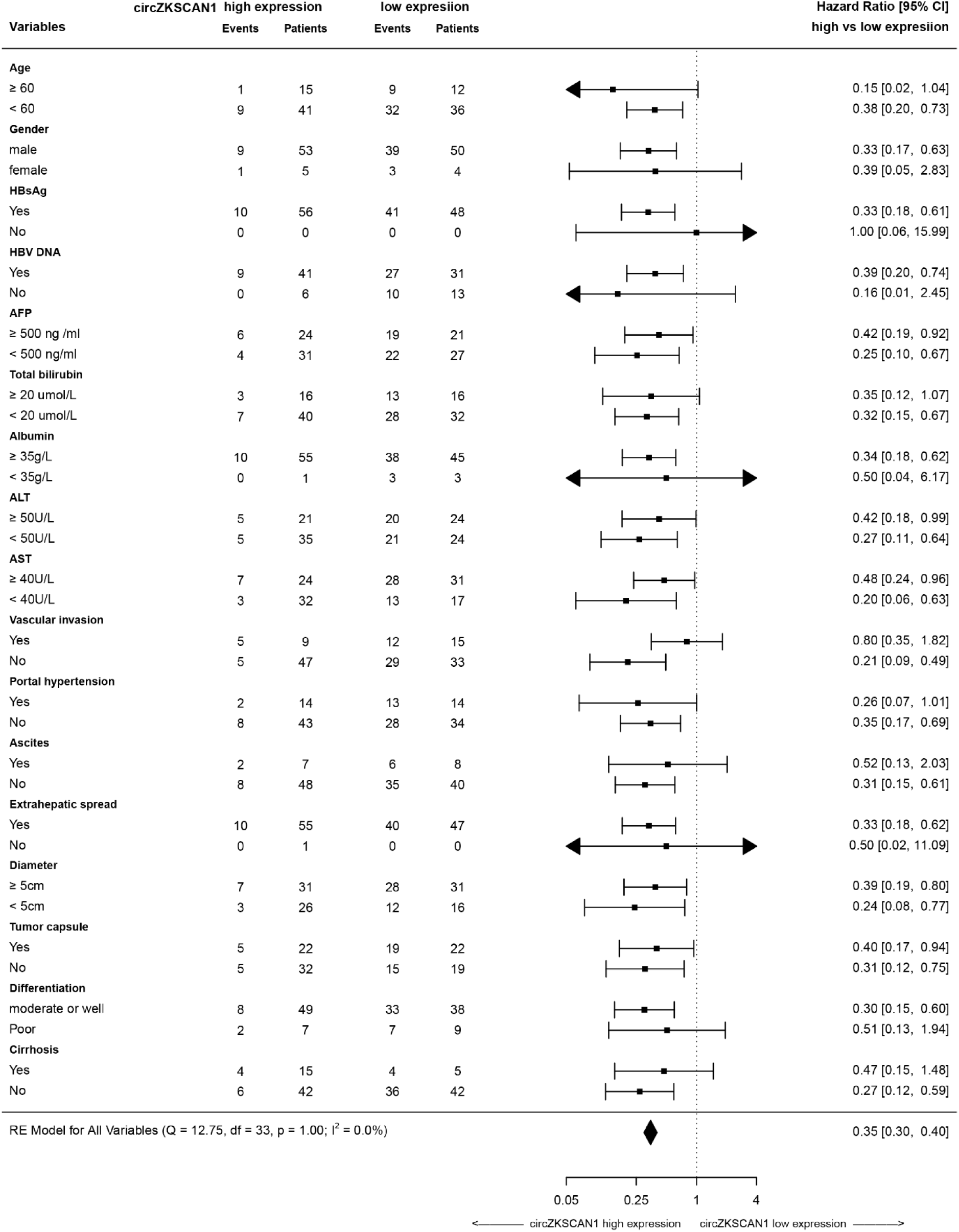
Forest plots of overall and progression-free survival in patient subgroups

**Figure 1.**
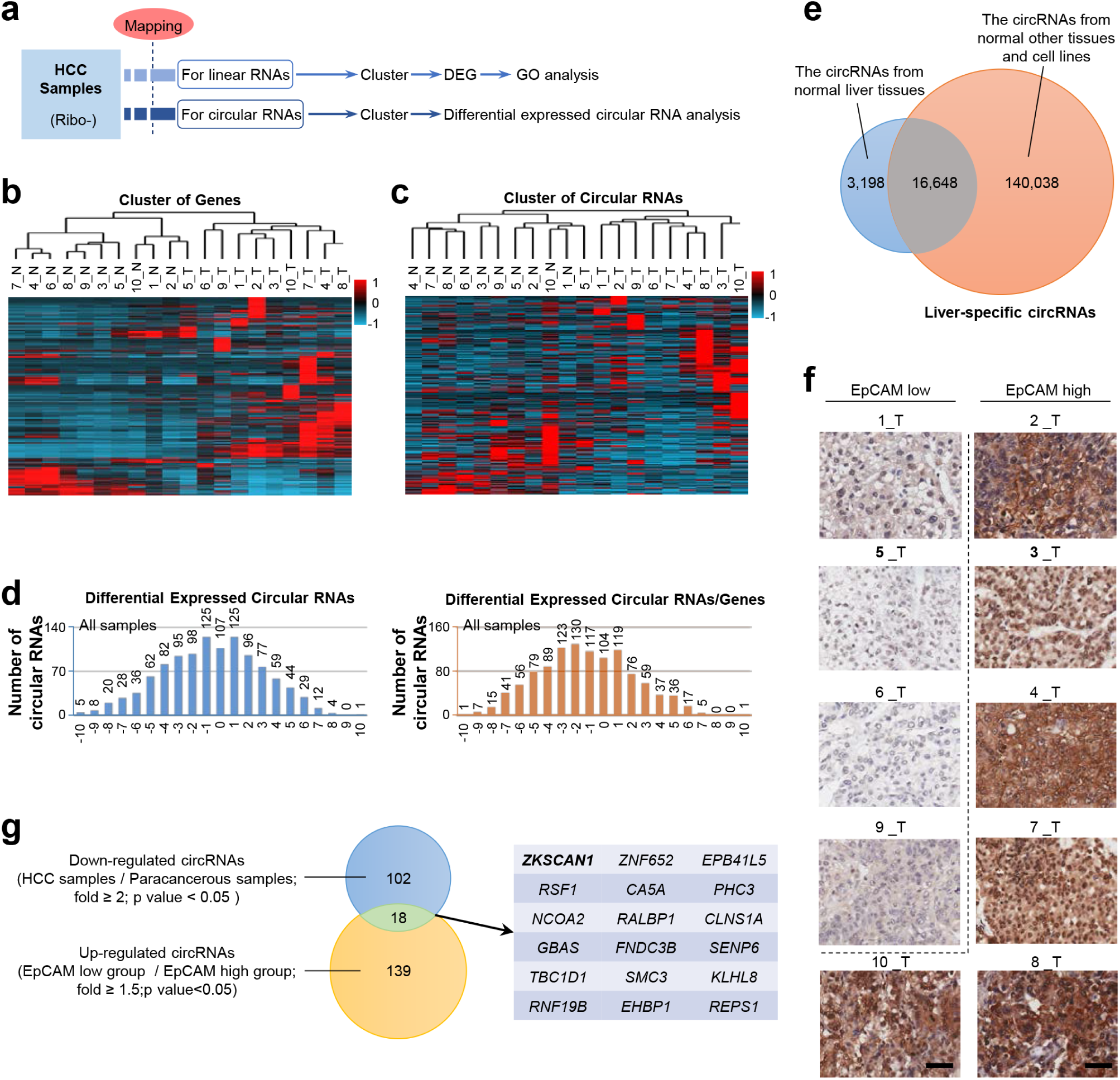
The exclusive circRNA expression spectrum in hepatocellular carcinoma. (a) Second-generation sequencing workflow. (b) Differentially expressed genes from an RNA-seq of 10 HCC samples and paired paracancerous tissues. n = 10, red is higher and blue is lower expression. (c) Circular RNAs cluster analysis from an RNA-seq of 10 HCC samples and paired paracancerous tissues. n = 10, red is higher and blue is lower expression. (d) Differentially expressed Circular RNAs in all cancerous tissue samples(left), as well as Circular RNAs that change independently after richness standardization with the corresponding mRNA(right). (e) Liver-specific circRNA spectrum. The overlap between circRNAs from normal liver tissues and normal other tissues and cell lines covers 16,648 liver-specific circRNAs. (f) Immunohistochemistry staining in formalin-fixed paraffin-embedded (FFPE) sections of 10 HCC sample. Scale bar, 200 µm. (g) Stemness-related circRNAs from sequencing. The overlap between Down-regulated circRNAs (HCC samples/Paracanceroussamples; fold >= 2;p value < 0.05) and Up-regulated circRNAs (EpCAM^low^group /EpCAM^high^group; fold >= 1.5;p value<0.05) covers 18 stemness-related circRNAs.

### 2. Specific features and prognostic value of circZKSCAN1 in HCC cells

After richness standardization of the original linear/circRNA sequencing data, we identified 120 circRNAs that were significantly downregulated in HCC. Among these, circZKSCAN1 expression was decreased > 2-fold in tumor tissues compared with paired non-tumor tissues. Interestingly, there was no change in expression levels of its linear counterpart, suggesting that circZKSCAN1 may function independently of its source gene ZKSCAN1 in HCC. Given the negative relationship between circZKSCAN1 and EpCAM, we investigated the underlying role of circZKSCAN1 in HCC CSCs. The human ZKSCAN1 gene is localized on human chromosome 17 and is characterized by a 2232 nucleotide-long region containing exons 2 and 3. Deep sequencing of various human cell lines revealed that the 3′ end of exon 3 and 5′ end of exon 2 could be spliced and connected to form a circRNA (Fig. 2a). To rule out the insertion of an intron between exons 2 and 3 during circularization, we designed two primers and subjected the PCR products to gel electrophoresis, which confirmed a direct connection between exons 2 and 3 (Supplementary Fig. 2a). Sequencing of the retrieved gel samples confirmed this conclusion (Supplementary Fig. 2b). The expression levels of circZKSCAN1 and linearZKSCAN1 detected in 120 HCC tissues and paired paracancerous tissues were in accordance with the previous findings (Fig. 2b). For further confirmation, we established a scoring system based on *in situ* hybridization staining intensity using a tissue microarray (n = 112; the baseline characteristics of the patients are shown in Supplementary Table 3) with the same results (Fig. 2c). Using the same 120 pairs of tissues, we identified a negative correlation between EpCAM mRNA and circZKSCAN1 expression (Fig. 2d). This negative correlation between EpCAM mRNA and circZKSCAN1 was confirmed using the *in situ* hybridization and immunohistochemistry scoring system (Fig. 2e). CircRNAs are known to be stable and relatively conserved. As expected, circZKSCAN1 was more resistant to RNase R treatment compared with its linear mRNA (Fig. 2f), supporting the existence of natural circZKSCAN1.

**Figure 2.**
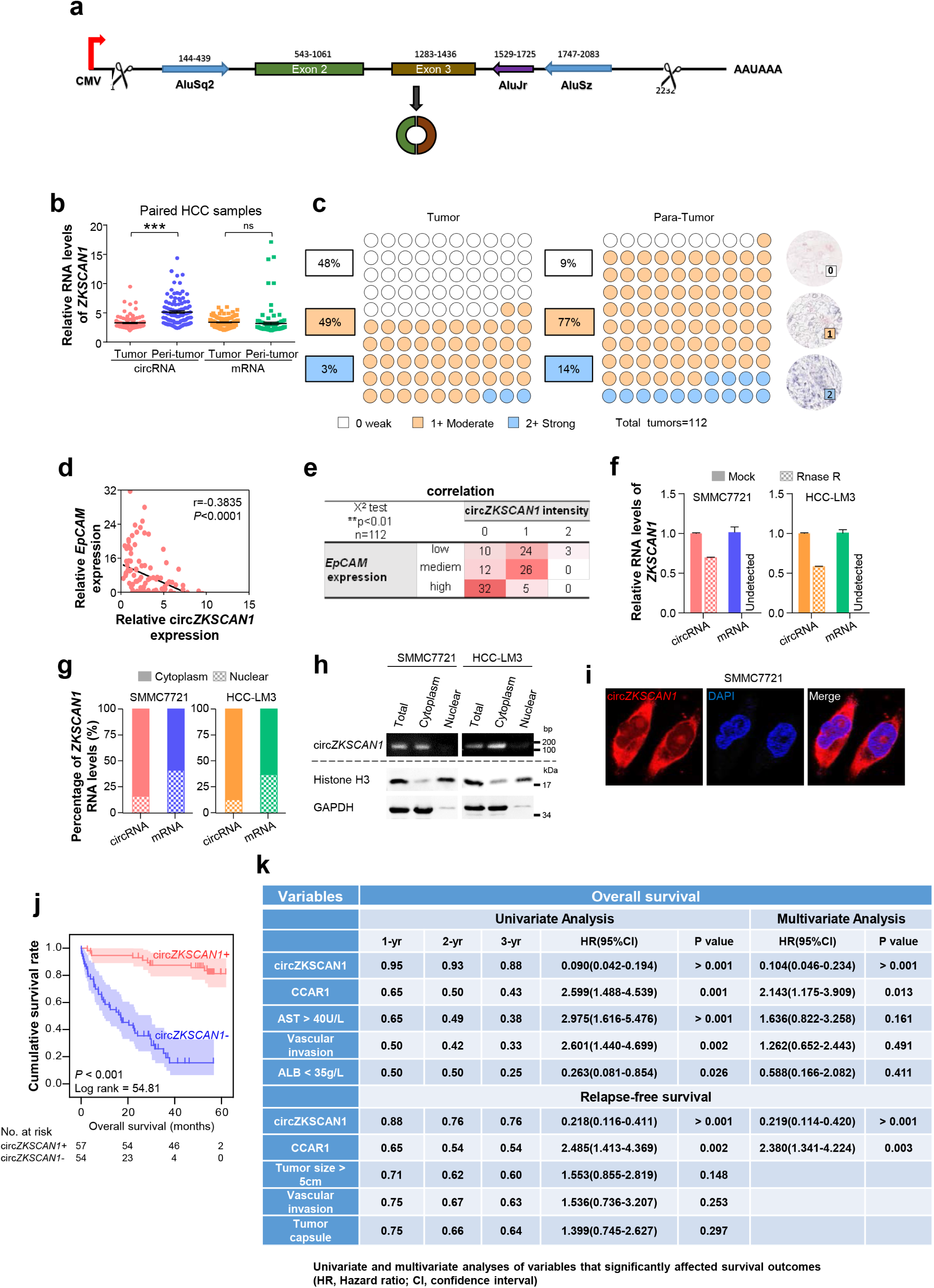
Specific features of circZKSCAN1 in HCC cells and its significant prognostic values. (a) Schematics of ZKSCAN1 circularization. A circular RNA is formed when the 5’ splice site at the end of exon 3 is joined to the 3’ splice site at the beginning of exon 2. Repetitive elements in the designated orientations and evolutionary conservation patterns are shown. (b) The relative expression of *ZKSCAN1* circRNA and mRNA were determined in fresh RNA samples of HCC tissue and paired non-tumor tissue (distal non-cancerous tissues away from HCC), n=112. (c) The expression level of circZKSCAN1 was measured by in situ hybridization staining using a pair of tissue microarrays (n=112). Score 0 defined as weak expression, score 1+ defined as moderate expression, score 2+ defined as strong expression. The staining intensity criteria was demonstrated on the right. The 100 dots shown in the figure represent the percentage not the exact number of tissues. (d) The correlation of EpCAM mRNA level and circZKSCAN1 expression level using 112 HCC frozen tissues was determined by qPCR. r=-0.3845, P<0.0001. (e) The correlation of EpCAM mRNA level and circZKSCAN1 ISH staining intensity. (f) Total RNAs were digested with RNase R followed by qRT-PCRs detection of circZKSCAN1expression. ZKSCAN1 mRNA was detected as the RNase R-sensitive control (n=4, Student t test). (g)The percent of *ZKSCAN1* levels. The expression of *ZKSCAN1* circRNA and mRNA were detected by qRT-PCRs in the nuclear and cytoplasm fractions of cells. (h) Immunoblot detection of cell cytoplasm and nucleus circZKSCAN1, GAPDH (cytoplasm control) and Histone3(nuclear control) of indicated cell lines are shown. (i) RNA-FISH assays were conducted to detect circZKSCAN1 expression in SMMC7721 cell using Cy3-labeled antisense probes(circZKSCAN1). Nuclei were stained with 4, 6-diamidino-2-phenylindole (DAPI). Scale bar, 10um. (j) Kaplan-Meier analysis of the correlation between circZKSCAN1 expression levels and overall survival (OS). (k) Univariate or multivariate analysis of HRs for overall survival and relapse-free survival. ***p<0.001; NS, not significant; HR, Hazard ratio; CI, confidence interval.

To clarify the function of circZKSCAN1, we identified its subcellular localization using a nucleus–cytoplasm separation technique. Unexpectedly, circZKSCAN1 was present almost exclusively in the cytoplasm, while linearZKSCAN1 was similarly distributed in the cytoplasm and nucleus (Fig. 2g and h). This was confirmed by fluorescence *in situ* hybridization (Fig. 2i). Based on the patient information for the tissue microarray (n = 112), we analyzed the data by correlation regression analysis and revealed that low circZKSCAN1 expression levels were associated with multiple HCC characteristics (Supplementary Table 4). Kaplan–Meier survival analysis showed that circZKSCAN1 expression level was positively correlated with overall and recurrence-free survival (Fig. 2j and Supplementary Fig. 2c). Univariate analysis of multiple survival and recurrence-related clinicopathological variables showed that circZKSCAN1, CCAR1 expression level, AST, ALB, and vascular invasion were significantly correlated with overall survival, and circZKSCAN1, CCAR1 expression level, tumor size (> 5 cm), vascular invasion, and tumor capsule were significantly correlated with recurrence-free survival. Each parameter was then subjected to multivariate analysis, which indicated that circZKSCAN1 and CCAR1 expression level were independent and significant factors affecting the survival of HCC patients (Fig. 2k). Furthermore, high circZKSCAN1 expression was associated with a favorable prognosis in most HCC patients, despite different characteristics (Table 1).

### 3. CircZKSCAN1 suppresses malignant behavior of HCC by downregulating cell stemness

We further characterized the role of circZKSCAN1 in regulating stemness in HCC using circZKSCAN1-knockdown and circZKSCAN1-overexpressing HCC cell lines, respectively (SMMC7721 and HCC-LM3), in *in vitro* and *in vivo* experiments. The efficiency of short hairpin RNA-mediated knockdown and lentivirus-mediated overexpression were confirmed by PCR (Fig. 3a and b). Expression levels of linear ZKSCAN1 were consistently unaffected by changes in expression of its circular form (Supplementary Fig. 3a).

**Figure 3.**
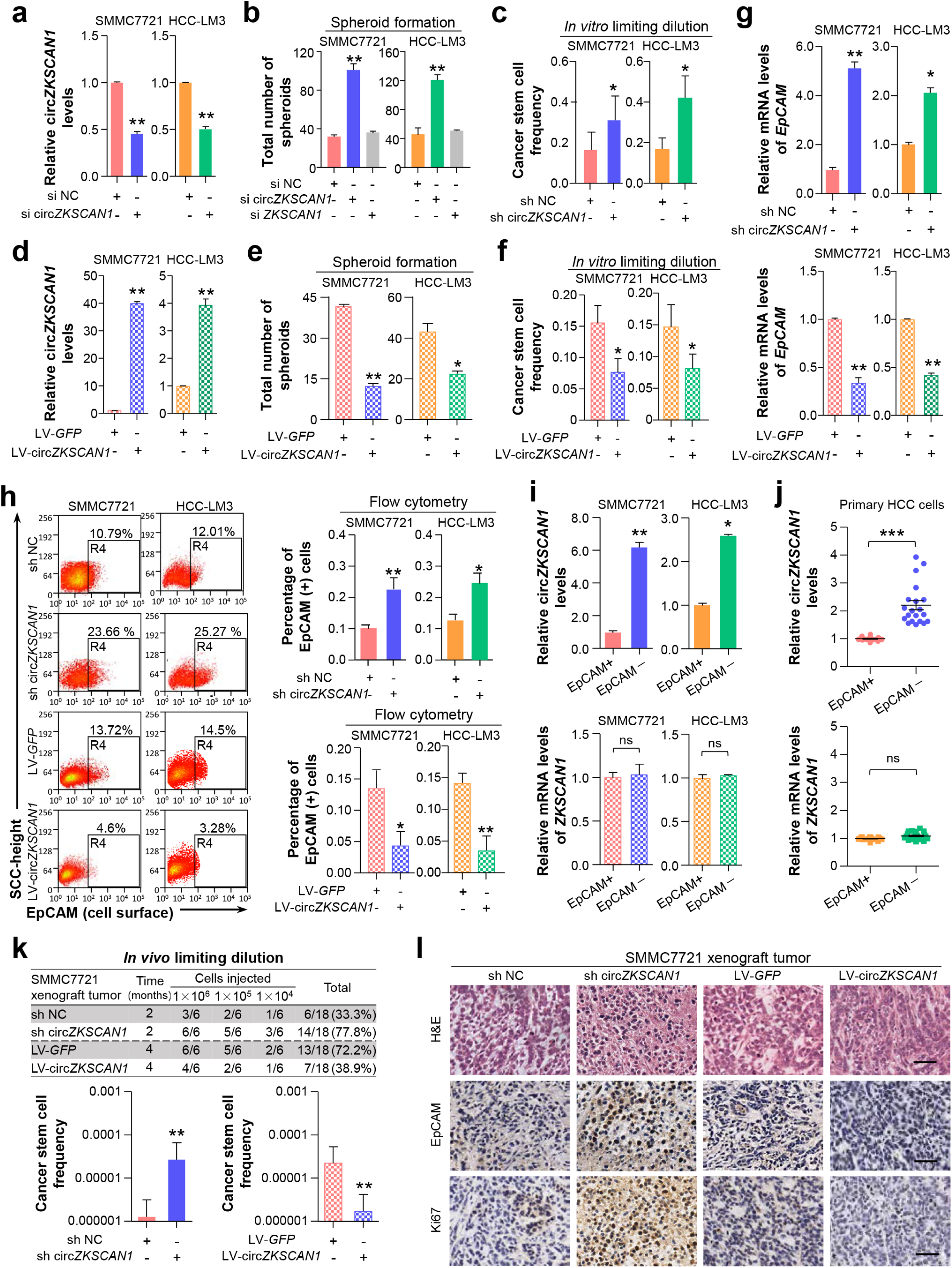
CircZKSCAN1 suppresses malignant behaviors of hepatocellular carcinoma by downregulating cellular stemness. (a) Expression of circZKSCAN1 was quantified by real-time PCR in SMMC7721/HCC-LM3 after transfected with two siRNAs against circZKSCAN1 or control siRNA (siNC), respectively. Data are presented as mean ± SEM (n=4). **P<0.01. (b) Spheroid formation assays were performed 48 hours after the indicated cell lines were treated with 100 M of circZKSCAN1 siRNA (si circZKSCAN1), ZKSCAN1 siRNA(si ZKSCAN1) or si NC. Results are shown as mean ± SEM, n=3. **P<0.01, based on the student’s t-test. (c)&(f) *In vitro* limiting dilution assays were performed to test cancer stem cell frequency of the indicated cell lines. Results are shown as mean ± SEM, n=3. *P<0.05. (d)The GFP-tagged circZKSCAN1 stably transfected SMMC7721/HCC-LM3 cell lines and their control cell lines were established using lentivirus. Expression of exogenous GFP-tagged circZKSCAN1 was quantified by real-time PCR. Data are presented as mean ± SEM (n=4). **P<0.01. (e)Spheroid formation assay were used to test the sphere formation abilities of the indicated cell lines. Results are shown as mean ± SEM, n=3. *P<0.05, **P<0.005, based on the student’s t-test. (g) The EpCAM mRNA expression level of different groups were confirmed by qPCR. Data are presented as mean ± SEM (n=4). *P<0.05, **P<0.01. (h) The EpCAM^+^population of different groups was determined by flow cytometry. SMMC7721(left); HCC-LM3(right). Data are presented as mean ± SEM (n=4). *P<0.05, **P<0.01. (i) The expression level of *ZKSCAN1* circRNA and mRNA in sorted EpCAM^+^and EpCAM^−^HCC cells. NS, not significant, *P<0.05, **P<0.01. (j) The expression level of *ZKSCAN1* circRNA and mRNA in sorted EpCAM^+^and EpCAM^−^primary HCC cells. NS, not significant, ***P<0.001. (k) Nude mice (n=6) were subcutaneously injected with the indicated number of SMMC7721 cells in which circZKSCAN1 was knocked-down or overexpressed using lentivirus before being implanted and the frequency of cancer stem cell was evaluated 2 or 4 months after injection by a limiting dilution assay. **p<0.01. (l) H&E and immunohistochemistry staining in formalin-fixed paraffin-embedded (FFPE) sections of SMMC7721 xenograft tumor. Scale bar, 200 µm.

We examined sphere formation and performed limiting dilution analysis *in vitro*. The sphere-formation ability of HCC cell lines was greatly enhanced by downregulation of circZKSCAN1 expression and markedly reduced by circZKSCAN1 overexpression (Fig. 3c and d). Intriguingly, numbers of CSCs increased after the circZKSCAN1 knockdown and showed improved colony-forming ability, while circZKSCAN1 overexpression had the opposite effects in HCC cell lines (Fig. 3e and f and Supplementary 3b, 3c). These results demonstrated that circZKSCAN1 was negatively correlated with cell stemness in HCC.

EpCAM is a distinctive CSC marker. We therefore subsequently explored the correlation between EpCAM and circZKSCAN1. CircZKSCAN1 suppressed the expression of EpCAM in modified HCC cell lines (Fig. 3g). Flow cytometry analysis supported this finding, with a higher frequency of EpCAM^+^cells among circZKSCAN1-knockdown cells, and vice versa (Fig. 3h). We separated HCC cells into EpCAM^+^and EpCAM^−^subgroups by magnetic bead sorting, and identified circZKSCAN1 downregulation among the EpCAM^+^subgroup, while its linear mRNA showed no similar trend (Fig. 3i).

Considering the potential differences between HCC cell lines and patient-derived HCC tumor cells, we also separated EpCAM^+/-^cells from 20 freshly resected HCC tumor masses from 10 patients by magnetic bead sorting. CircZKSCAN1 was greatly downregulated in patient-derived EpCAM^+^HCC cells, while linearZKSCAN1 showed no such difference (Fig. 3j).

We investigated the role of circZKSCAN1 *in vivo* using mice injected subcutaneously with modified HCC cell lines. Tumor formation was increased following subcutaneous injection of as few as 10,000 circZKSCAN1-knockdown HCC cells (Fig. 3k). Furthermore, tumors in the circZKSCAN1-knockdown group were significantly larger (Supplementary Fig. 3e). We examined circZKSCAN1 expression levels in the xenografts by PCR (Supplementary Fig. 3d). CircZKSCAN1-overexpressing HCC cell lines showed reduced tumor-forming ability, with a lower incidence of tumors and smaller tumor size (Fig. 3k and Supplementary Fig. 3f). We examined the correlation between the number of EpCAM^+^cells and circZKSCAN1 expression in xenograft tumors in mice by immunohistochemical staining (Supplementary Fig. 3d, 3f). The numbers of EpCAM^+^cells were increased in circZKSCAN1-knockdown and decreased in circZKSCAN1-overexpressing tumors (Fig. 3i).

We also determined if circZKSCAN1 expression affected the proliferative ability of HCC. Interestingly, transient silencing of circZKSCAN1 facilitated the proliferation of HCC cell lines (Huh-7, HCC-LM3, and SMMC7721) and a normal hepatocyte cell line (QSG7701), further indicating a close connection between circZKSCAN1 and malignant behavior in HCC (Supplementary Fig. 3g, 3h). In contrast, silencing linear ZKSCAN1 had no effect on the proliferation of either HCC or normal hepatocyte cell lines (Supplementary Fig. 3i, 3j). A similar result was observed in constructed cell lines (Supplementary Fig. 3k, 3l).

Silencing circZKSCAN1 also enhanced the metastatic abilities of both normal hepatocytes and HCC cells as shown by cell migration assay (data not shown).

### 4. CircZKSCAN1 acts by sponging specific RBPs

CircRNAs mostly act by competing against miRNAs and RBPs to regulate their target genes. We performed bioinformatics analysis and identified 10 RBPs (FMRP, FUS, DGCR8, ELAVL, EIF4A3, PTB, IGF2BP1, IGF2BP2, LIN28A, LIN28B) that may compete against circZKSCAN1 (14). Knockdown of any single RBP did not affect spheroid formation in HCC control cells, but knockdown of FMRP significantly compromised spheroid-formation ability in circZKSCAN1-deficient HCC cell lines (Fig. 4a). Unexpectedly, double knockdown of both circZKSCAN1 and FMRP could ameliorate the enhanced stemness of both SMMC7721 and HCC-LM3 cell lines induced by single knockdown of circZKSCAN1, demonstrating that circZKSCAN1 might suppress stemness by confining the biological functions of FMRP (Fig. 4b, 4c). Flow cytometry analysis also showed that double knockdown of circZKSCAN1 and FMRP could restore EpCAM^+^cell numbers upregulated by circZKSCAN1 deficiency (Fig. 4d). Double knockdown of circZKSCAN1 and FMRP could counteract the improvement effect on cellular proliferation induced by knockdown of circZKSCAN1 alone (Supplementary Fig. 4a).

**Figure 4.**
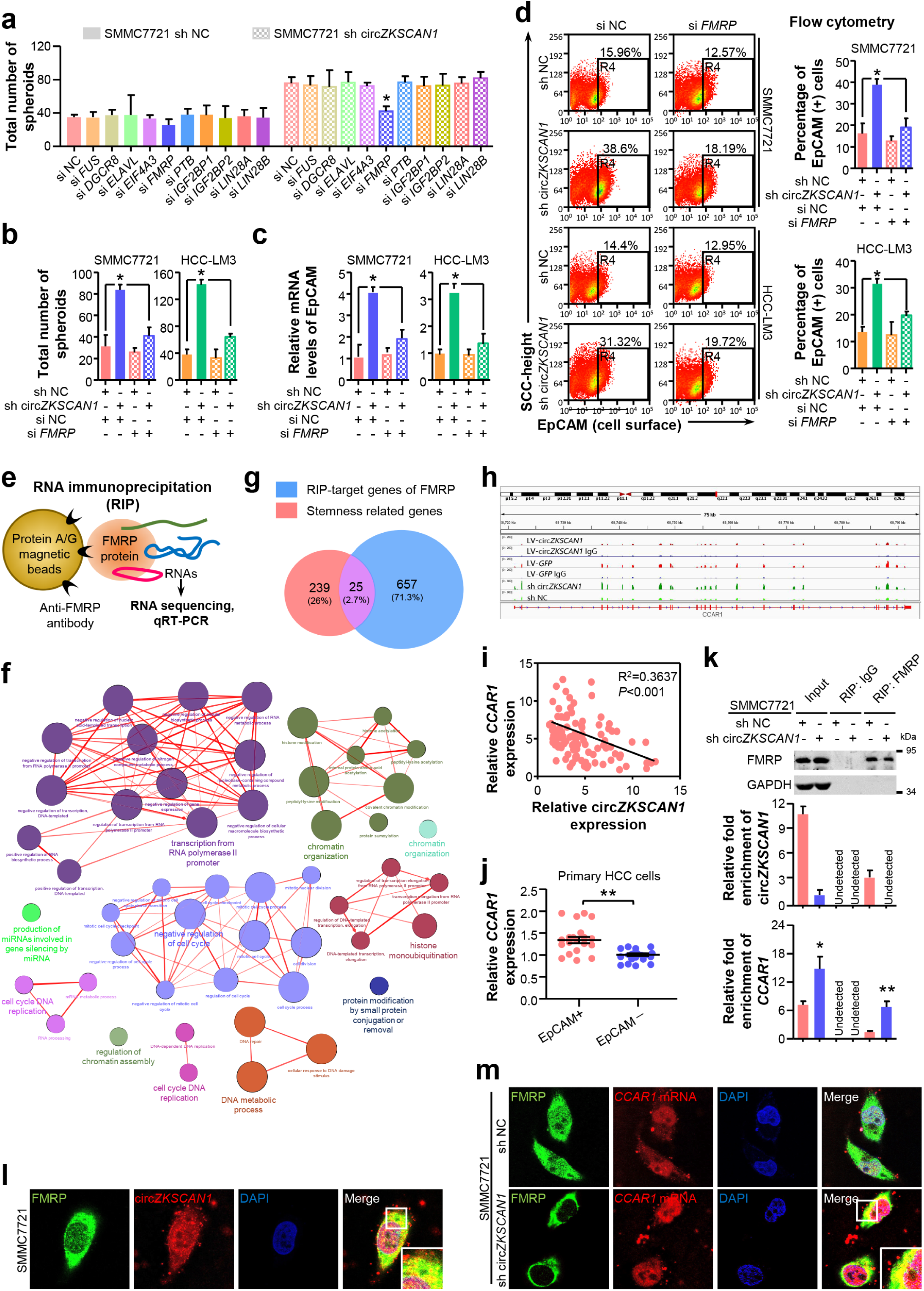
CircZKSCAN1 performs its function through sponging specific RNA-binding protein. (a) Spheroid formation assay result in the context of 10 different RBP knockdown. HCC control cell line (left); HCC lentivirus-mediated circZKSCAN1 deficient cell line (right). Data are presented as mean ± SEM (n=4). *P<0.05. (b) Comparison of the spheroid formation ability between single circZKSCAN1 or FMRP knockdown and double knockdown. Data are presented as mean ± SEM (n=4). *P<0.05. (c) The EpCAM mRNA expression level of different groups were confirmed by qPCR. Data are presented as mean ± SEM (n=4). *P<0.05. (d) The EpCAM^+^population of different groups was determined by flow cytometry. SMMC7721(above); HCC-LM3(below). Data are presented as mean ± SEM (n=4). *P<0.05. (e) Graphical abstract of the RNA immunoprecipitation sequencing procedure. (f) GO analysis of the RIP-target genes of FMRP. (g) The overlap between RIP-target genes of FMRP and stemness-related genes covers 25 genes. (h) RIP-seq results of FMRP binding sites in CCAR1 gene. (i) The correlation of CCAR1 mRNA level and circZKSCAN1 expression level using 120 HCC frozen tissues was determined by qPCR. R^2^=0.3637, P<0.001. (j) The expression level of CCAR1 in sorted EpCAM^+^and EpCAM^−^primary HCC cells. P<0.001. (k) Whole-cell RNA extract was precipitated with input (positive control), IgG (negative control) and FMRP antibody and the precipitation was then verified by Western Blot. CircZKSCAN1 and CCAR1 level of the precipitations was confirmed by qPCR. (l) The co-localization of FMRP protein and circZKSCAN1 was confirmed by immunofluorescence and fluorescence in situ hybridization. (m) The co-localization of FMRP protein and CCAR1 was confirmed by immunofluorescence and fluorescence in situ hybridization. Sh NC, HCC control cells. Sh circZKSCAN1, HCC lentivirus-mediated circZKSCAN1 knockdown cells.

We investigated the potential target stemness-related genes of FMRP by RNA immunoprecipitation (RIP)-sequencing (Fig. 4e). Mapping data revealed that FMRP had 682 potential target genes, and further GO analysis demonstrated that these target genes had broad functions in cellular physiological processes (Fig. 4f). We examined the overlap between these 682 genes and a stemness-related gene signature containing 264 genes, and selected 25 genes identified in both (Fig. 4g). Based on previous studies, we narrowed the list down to six genes with documented roles in cancers: CCAR1 (15), ARHGAP21 (16), PPP4R1 (17), ZHX3 (18), GOLGA8B, and ANKRD18B (19). Further correlation analysis demonstrated that of these six genes, CCAR1 was negatively correlated with circZKSCAN1 (Fig. 4h, 4i, and Supplementary Fig. 4b). Using DNA extracted from 10 freshly resected HCC tissues, as noted above, we confirmed that CCAR1 expression levels were significantly higher in sorted EpCAM^+^HCC cells, suggesting that CCAR1 had a positive effect on stemness regulation (Fig. 4j). CircZKSCAN1 deficiency had no effect on the expression level of FMRP, but markedly boosted the expression of CCAR1, as shown by RIP (Fig. 4k and Supplementary Fig. 4c). Considering the positive correlation between CCAR1 and EpCAM, we confirmed that circZKSCAN1 suppressed CSCs by regulating the expression of CCAR1.

We carried out fluorescence *in situ* hybridization at a subcellular level and determined that FMRP was localized exclusively in the cytoplasm, consistent with The Human Protein Atlas, and co-localization of FMRP protein and circZKSCAN1 was confirmed (Fig. 4l). We also demonstrated direct binding between FMRP protein and CCAR1 mRNA (Fig. 4m).

### 5. FMRP modulates cellular stemness in HCC through its target gene CCAR1, which assists in the activation of the Wnt/β-catenin pathway

CircZKSCAN1 had a negative regulatory effect on CCAR1 expression, as demonstrated by RIP (Fig. 4k). This negative correlation between circZKSCAN1 and CCAR1 was confirmed in circZKSCAN1-modified HCC cells at both the protein and mRNA levels (Fig. 5a). We therefore further explored the role of CCAR1 in cell stemness regulation. We transfected HCC cells with small interfering RNA (siRNA) targeting CCAR1 and confirmed the knockdown efficiency by RT-PCR (Supplementary Fig. 5a). We then explored the importance of CCAR1 in the regulation of stemness by circZKSCAN1 by *in vitro* sphere-formation assay in the setting of CCAR1 deficiency or combined CCAR1 and circZKSCAN1 deficiency. The loss of CCAR1 could counteract the enhancement of stemness induced by circZKSCAN1 deficiency, suggesting that CCAR1 and circZKSCAN1 had opposite effects on stemness (Fig. 5b). Furthermore, knockdown of circZKSCAN1 boosted the expression of CCAR1, while changes in the expression of CCAR1 had no effect on circZKSCAN1, suggesting that the circZKSCAN1–FMRP–CCAR1 axis might not involve positive or negative feedback (Supplementary Fig. 5b). Moreover, flow cytometry indicated that EpCAM expression was inhibited in CCAR1-knockdown HCC cells (Fig. 5c), consistent with the above findings (Fig. 4j). CCAR1 was previously reported to act as a co-activator dependent on β-catenin function in the Wnt signaling pathway (20). Consistently, we confirmed that CCAR1 had no effect on β-catenin nuclear translocation (Supplementary Fig. 5c).

**Figure 5.**
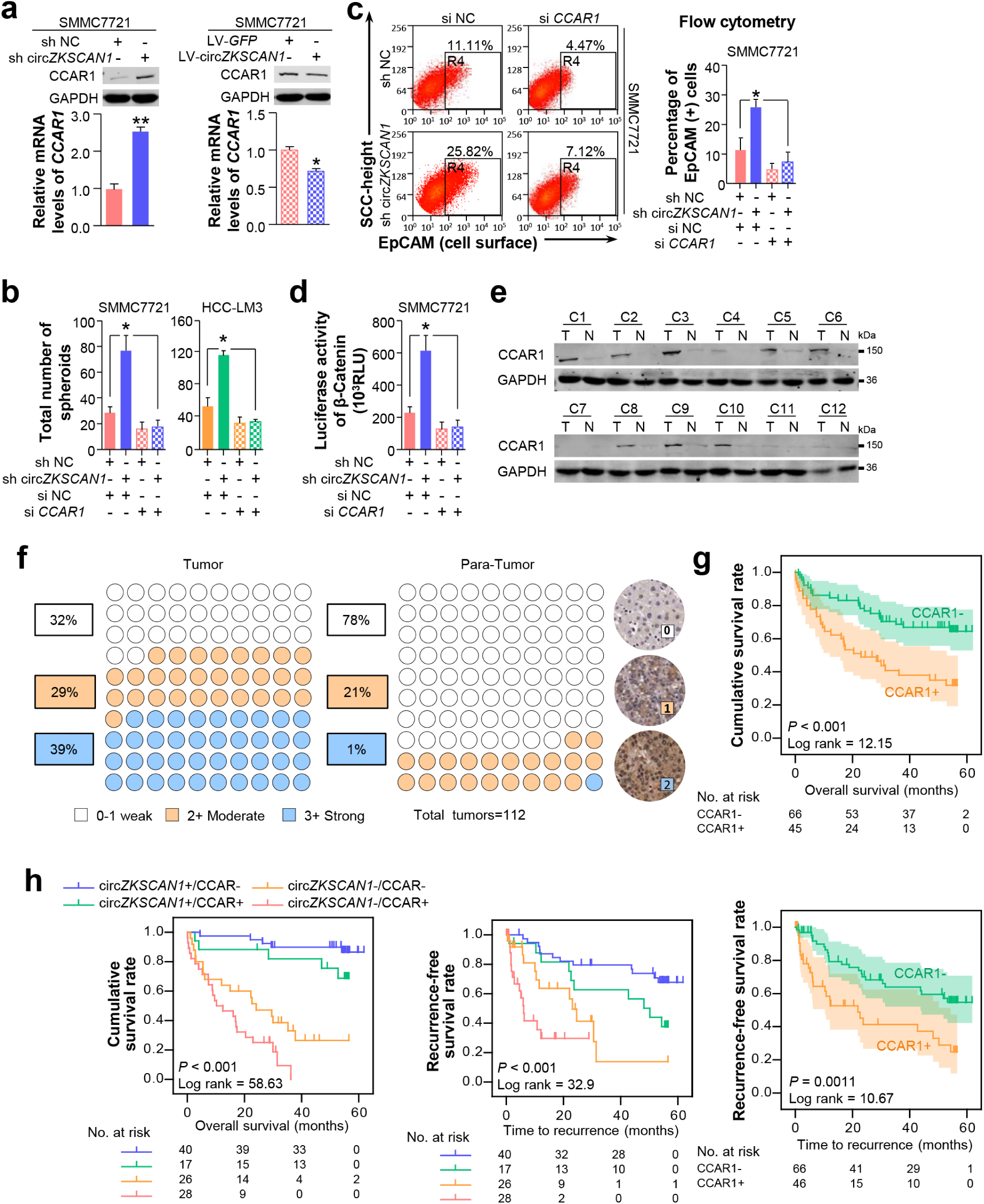
FMRP modulates cellular stemeness in HCC through its target gene, CCAR1 which assists in the activation of Wnt/β-catenin pathway. (a) The expression level of CCAR1 protein and mRNA in stable constructed circZKSCAN1 knockdown or overexpression HCC cell line were determined by Western Blot or qPCR. Data are presented as mean ± SEM (n=4). *P<0.05; \*\**P*<0.01. (b) Comparison of the spheroid formation ability between single circZKSCAN1 or CCAR1 knockdown and double knockdown. Data are presented as mean ± SEM (n=4). *P<0.05. (c) The EpCAM^+^population of different groups was determined by flow cytometry. Data are presented as mean ± SEM (n=4). *P<0.05. (d) The transcriptional activity of β-catenin downstream targets was measured by luciferase reporter gene assay. Data are presented as mean ± SEM (n=4). *P<0.05. (e) The expression level of CCAR1 protein was measured in 12 HCC tissues and paired paracancerous tissues. (f) The expression level of CCAR1 was measured by immunohistochemistry staining using a pair of tissue microarrays (n=112). Score 0-1 defined as weak expression, score 2 defined as moderate expression, score 3 defined as strong expression. The staining intensity criteria was demonstrated on the right. The 100 dots shown in the figure represent the percentage not the exact number of tissues. (g) Kaplan-Meier analysis of the correlation between CCAR1 expression levels and overall survival (OS) or recurrence free survival (RFS) (h) The prognostic value of the combination of circZKSCAN1 and CCAR1 was compared with individual parameter.

We then evaluated the transcriptional activity of T cell factor/lymphoid enhancer binding factor 1 by luciferase analysis using pGL3-OT and pGL3-OF which showed that knockdown of CCAR1 ameliorated the transcriptional activity of Wnt signaling downstream genes (Fig. 5d).

Notably, CCAR1 expression was markedly upregulated in HCC compared with paracancerous tissues (Fig. 5e), thus establishing it as a crucial oncogene in HCC. We further determined the expression pattern of CCAR1 in HCC via immunohistochemical staining using a tissue microarray consisting of paired cancerous and paracancerous tissues from 112 patients. As expected, CCAR1 was significantly enriched in HCC compared with normal tissues (39% versus 1% strong staining, respectively) (Fig. 5f). We also analyzed the prognoses of these 112 patients which showed that CCAR1 expression level was negatively correlated with survival (Fig. 5g), thus supporting the role of CCAR1 as a critical oncogene. The previous multivariate analysis also identified CCAR1 as an independent significant factor in OS (overall survival) and RFS (recurrence-free survival) in HCC patients (Fig. 2k). Furthermore, the combination of circZKSCAN1 and CCAR1 showed significant prognostic value for predicting overall and recurrence-free survival in HCC patients, with improved predictive power compared with either parameter alone (Fig. 5h).

### 6. Overexpression of circZKSCAN1 directly inhibits HCC growth *in vivo* which was regulated by QKI

Given that circZKSCAN1 acts as a tumor suppressor in HCC, its therapeutic potential should be evaluated further. We, therefore, constructed four PDX (patient-derived xenograft) mouse models (derived from the above 20 freshly resected HCC tissues). The mice were injected with circZKSCAN1-overexpressing adenovirus into the tumor mass (Fig. 6a), and the tumors were harvested 18 days after injection. CircZKSCAN1 overexpression significantly reduced the growth rate of the tumor xenografts compared with the control group (Fig. 6b, 6c). CircZKSCAN1 expression levels in both groups were confirmed by RT-PCR (Fig. 6d). Moreover, we confirmed that ZKSCAN1 mRNA levels were similar in both groups, while CCAR1 was markedly upregulated in the circZKSCAN1-overexpressing group (Fig. 6d).

**Figure 6.**
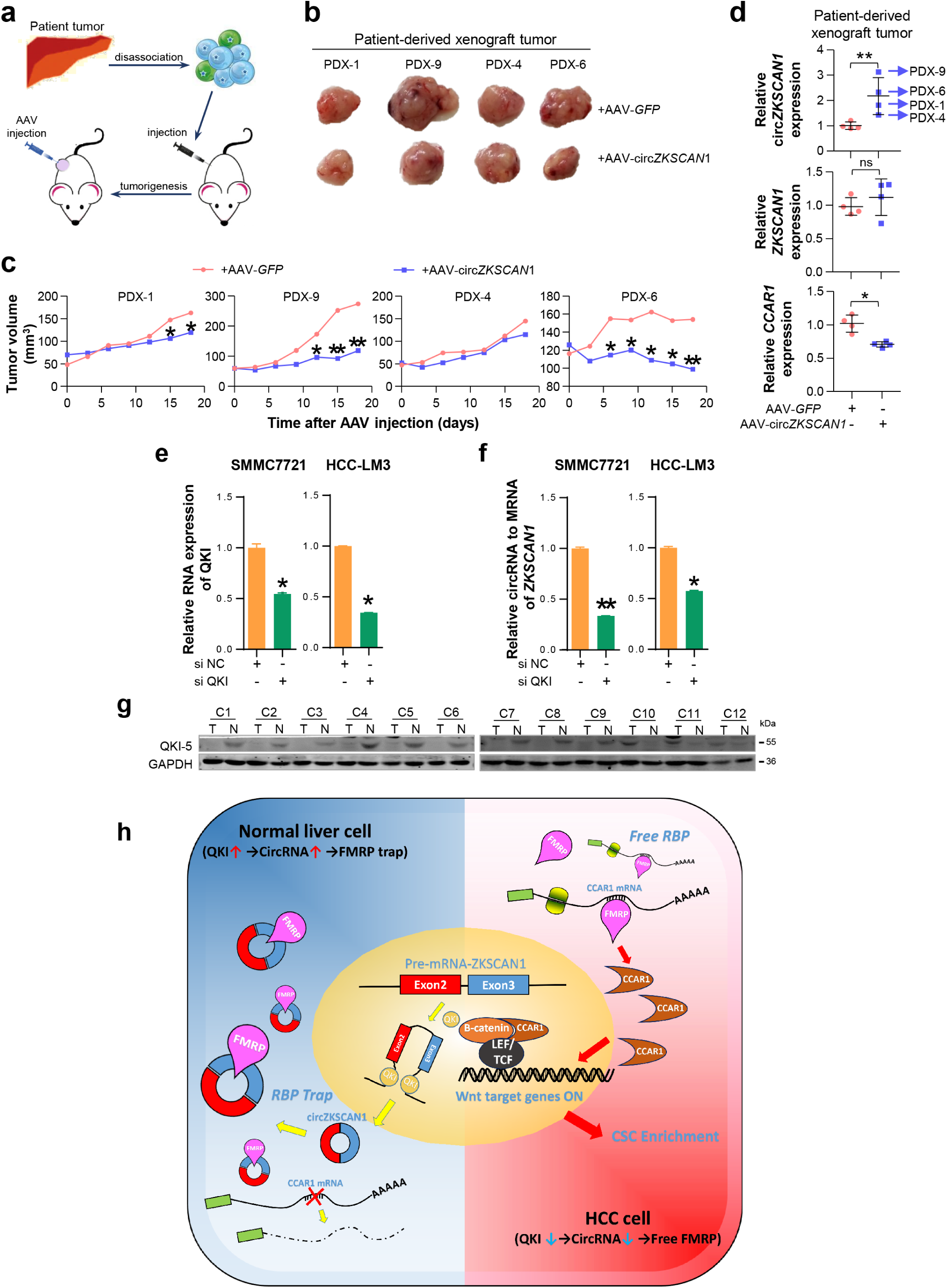
The overexpression of circZKSCAN1 directly inhibits HCC growth in vivo and the regulation of circZKSCAN1. (a) Graphical abstract of the intratumoral injection. (b) Harvested tumor mass from 4 PDX models were presented. Control group (above), CircZKSCAN1 overexpression group (below). (c) The volume of PDX tumors were measured every 3 days. Growth curves were presented. (d) The expression level of circZKSCAN1, ZKSCAN1 mRNA and CCAR1 mRNA of the PDX tumors (n=4) were determined by qPCR. The different degrees of significance were indicated as follows in the graphs: \**P*<0.05; **P<0.01. (e) The knockdown efficiency of small interfering RNA targeting pan-QKI was determined by qPCR. Data are presented as mean ± SEM (n=4). The different degrees of significance were indicated as follows in the graphs: \**P*<0.05. (f) The expression level of standardized circZKSCAN1 in the context of pan-QKI deficiency was determined by qPCR. Data are presented as mean ± SEM (n=4). *P<0.05; **P<0.01. (g) The expression level of QKI-5 protein in 12 pairs of tumor and para-tumor tissues was measured by Western Blot. (h) Graphical abstract of the whole QKI5-circZKSCAN1-FMRP-CCAR1 axis.

We then investigated the abnormal downregulation of circZKSCAN1 in HCC. The Quaking (QKI) protein family was previously reported to play an important role in RNA circularization, and QKI5 was shown to regulate circZKSCAN1 formation (21). We knocked down QK1 expression using a targeted siRNA and confirmed the knockdown efficiency by PCR (Fig. 6e). QKI knockdown significantly affected the expression of circZKSCAN1, consistent with the previous findings (Fig. 6f). Besides, the expression level of QKI protein was downregulated in HCC tissue which was confirmed by Western Blot (Fig. 6g). Overall, we concluded that abnormal downregulation of QKI in HCC tissues contributed to the reduced expression of circZKSCAN1.

## Discussion

CircRNAs were first discovered in RNA viruses (22) but received little attention because of their apparently low levels in cells. However, second-generation sequencing techniques and bioinformatics have revealed that some circRNAs are much more abundant than the mRNAs from which they originate. CircRNAs are now widely recognized as a novel subset of competing endogenous RNAs that regulate target genes by affecting the functions of miRNAs and RBPs. The roles of circRNAs have been studied in various cancers, including liver (11), lung (23), breast (24), bladder (25), and gastrointestinal cancer (26), and have been shown to facilitate proliferation, metastasis, and chemotherapeutic resistance (23,27). However, to the best of our knowledge, few studies have reported on the possible effects of circRNAs on CSCs. CSCs have been shown to be responsible for tumor initiation, progression, metastasis, chemoresistance, and relapse (28). The results of the current study provide the first evidence to show that circZKSCAN1 suppresses cell stemness in HCC by competing against the RBP, FMRP.

We identified possible circRNAs with regulatory functions in HCC stem cells using paired tumor and paracancerous tissues from 20 HCC patients, separated into EpCAM^high^(n = 6) and EpCAM^low^(n = 4) groups based on immunohistochemical staining scores. We then acquired the complete expression profile by next-generation sequencing of all samples and successfully filtered out 157 circRNAs significantly enriched in the EpCAM^low^group using real-time PCR and siRNAs targeting specific circRNAs. After comparing the expression levels of these circRNAs and their corresponding mRNAs in cancerous and paracancerous tissues, we selected circZKSCAN1, which was expressed independently of its mRNA, as a circRNA with a potentially unique role in HCC. Further experiments confirmed that circZKSCAN1 inhibited cell stemness, proliferation, and metastasis in HCC.

Although the relationship between circRNAs and cancers has been widely studied, the underlying mechanisms remain largely obscure. The best-known function of circRNAs in cancers is to act as a miRNA sponge and compete for mRNA-binding sites on miRNAs (6,8). For example, circRNA hsa_circ_000984 can facilitate the growth and metastasis of colon cancer by sponging miR-106b (29). However, the ability of circRNAs to modulate malignant behavior via RBPs has not previously been reported. RBPs play a major role in the post-transcriptional control of RNAs, such as in splicing, polyadenylation, and in mRNA stabilization, localization, and translation. RBPs are therefore likely to play a part in the regulation of circRNAs in cancers. The results of this study extend our understanding of their role by demonstrating that circZKSCAN1 inhibited stemness in HCC by acting as an RBP (FMRP) sponge, rather than a miRNA sponge.

FMRP mainly functions in the nervous system, where it acts as an RBP and controls the translation of its target mRNAs (30). FMRP1 gene knockdown causes the relatively common genetic intellectual disability, fragile X syndrome (31). Although FMRP has mostly been studied in the field of neuroscience, its expression is ubiquitous, suggesting that it may also play important roles in other organs and diseases, such as cancer (30). Previous studies showed that FMRP expression was upregulated in HCC cells (32). Furthermore, the risk of cancer was shown to be significantly lower in patients with fragile X syndrome (33). However, the precise regulatory mechanism of FMRP in cancer biology remains unknown. The current results suggest that circZKSCAN1 and FMRP act to control HCC cell stemness in opposite directions, such that increased stemness induced by circZKSCAN1 knockout can be ameliorated by FMRP deficiency. We also demonstrated that FMRP enhanced stemness in HCC cells by regulating the expression of CCAR.

CCAR1 plays a crucial role in cell proliferation and apoptosis pathways and was originally identified as a perinuclear phosphoprotein triggering apoptosis signaling in breast cancer in a retinoid-dependent manner (34). Other studies reported that CCAR1 participated in cell proliferation and apoptosis in various cell lines, including cancer cells (35,36). The Wnt/β-catenin signaling pathway is a key pathway involved in CSC regulation. Furthermore, CCAR1 has been confirmed as a functional β-catenin-binding protein, which binds directly with lymphoid enhancer binding factor 3/T cell factor family members and supports β-catenin in the transcriptional activation of Wnt target genes, thus further enhancing cell stemness in colon cancer (20). Based on this information, we performed gain-and loss-of-function experiments *in vitro* and demonstrated that CCAR1 was positively correlated with cell stemness in HCC through upregulating levels of active β-catenin. Interestingly, CCAR1 was expressed almost exclusively in HCC compared with normal liver tissues, based on 20 sets of paired cancerous and paracancerous tissues, suggesting that CCAR1 may play a more complex role in hepatocarcinogenesis, in addition to regulating cell stemness. Ha et al. also reported that overexpression of CCAR1 in HCC was associated with a poor prognosis (37).

Despite the fact that several circRNAs have been shown to be dysregulated in cancers, there is currently no evidence for the mechanisms underlying this phenomenon. Conn et al. discovered that the RBP and alternative splicing factor QKI5 regulated the formation of thousands of circRNAs, including circZKSCAN1, during epithelial-mesenchymal transition (21). We further identified QKI5 as significantly downregulated in HCC tissues compared with paired paracancerous tissues, leading to the differential expression of circZKSCAN1, as confirmed in our further experiments. QKI5 may thus act as a tumor suppressor in HCC. However, previous studies demonstrated that QKI protein may play dual roles in cancers, by promoting proliferation and invasion in colon cancer but inhibiting tumor progression in prostate cancer (38,39). QKI itself might thus function as an oncogene in HCC, independent of the function of circZKSCAN1. Further experiments are needed to clarify the relationship between QKI and HCC.

Although most circRNAs are classified as non-coding RNAs, some possess coding potential (40). However, the coding potential of circZKSCAN1 was not considered in the current study and should thus be addressed in future research.

Overall (Fig. 6h), this study demonstrated that circZKSCAN1 is downregulated by QKI in HCC and inhibits multiple malignant behaviors by suppressing cell stemness. Mechanistically, circZKSCAN1 acts as an RBP sponge, competing against the FMRP target gene, CCAR1, and subsequently deactivating the Wnt/β-catenin signaling pathway. Furthermore, circZKSCAN1 induced tumor quiescence *in vivo*, thus indicating its potential for precision targeted treatment. These results thus provide novel insights into the potential role of the QKI5– circZKSCAN–FMRP–CCAR1 axis in the therapeutic management of HCC.

**Author Contributions**
Yan-Jing Zhu, Bo Zheng, Xu-Kai Ma and Xin-Yuan Lu performed all the experiments. Xi-Meng Lin, Gui-Juan Luo and Shuai Yang provided human specimens, clinical information and data analysis. Qing Zhao, Xin Chen, Ying-Cheng Yang, Xiao-Long Liu, Rui Wu, Jing-Feng Liu, Yang Ge and Li Yang provided support with experimental materials and techniques. Hong-Yang Wang and Lei Chen designed research and wrote the manuscript.

## Abbreviations

HCC: hepatocellular carcinoma
circRNA: Circular RNA
CCAR1: cell cycle and apoptosis regulator 1
FMRP: fragile X mental retardation protein
CSC: cancer stem cells
EpCAM: epithelial cell adhesion molecule
RBPs: RNA-binding proteins
ZKSCAN1: Zinc-finger protein with KRAB and SCAN domains 1.

